# Mesenchymal Stromal Cell secretome is affected by tissue source, donor age and sex

**DOI:** 10.1101/2023.01.30.526247

**Authors:** AJ Turlo, DE Hammond, KA Ramsbottom, J Soul, A Gillen, K McDonald, MJ Peffers, PD Clegg

**Affiliations:** Department of Musculoskeletal and Ageing Science, Institute of Life Course and Medical Sciences, University of Liverpool; Department of Cellular and Molecular Physiology, Institute of Systems, Molecular and Integrative Biology, University of Liverpool; Computational Biology Facility, Liverpool Shared Research Facilities, Faculty of Health and Life Sciences, University of Liverpool; Department of Veterinary Science, Philip Leverhulme Equine Hospital, University of Liverpool; Biobest Laboratories Ltd

**Keywords:** mesenchymal stromal cells, secretome, proteomics, age, equine, donor

## Abstract

Variation in Mesenchymal Stromal Cell (MSC) function depending on their origin is problematic, as it may confound clinical outcomes of MSC therapy. Current evidence suggests that the therapeutic benefits of MSCs is primarily attributed to secretion of various biologically active factors (secretome). However, the effect of donor characteristics on the MSC secretome composition remains largely unknown. Here, we examined the influence of donor age, sex and tissue source, on the protein profile of the equine MSC secretome. Initially, we used dynamic metabolic labelling with stable isotopes combined with liquid chromatography-tandem mass spectrometry (LC-MS/MS) to identify secreted proteins in MSC conditioned media (CM). Seventy proteins were classified as classically-secreted based on the rate of isotope label incorporation into newly synthesised proteins released into the extracellular space. Next, we analysed CM of bone marrow- (n = 14) and adipose-derived MSCs (n = 16) with label-free LC-MS/MS proteomics. Clustering analysis of 314 proteins detected across all samples identified tissue source as the main factor driving variability in MSC CM proteomes. Linear modelling applied to the subset of 70 secreted proteins identified tissue-related difference in the abundance of 23 proteins. There was an age-related decrease in the abundance of two proteins (CTHRC1, LOX), which has been validated with western blot and enzymatic activity assay. There was limited evidence of sex-related differences in protein abundance. In conclusion, this study provides evidence that tissue source and donor age contribute to heterogeneity in the protein composition of MSC secretomes which may influence the effects of MSC-based cell therapy.

**Significance statement:** This research shows for the first time that donor age can influence the proteins secreted by mesenchymal stromal cells (MSC), which is considered the main mechanism through which they mediate their biological actions. This information may improve our understanding of the variable outcomes observed in MSC clinical studies, as well as identify optimal sources and donor selection for specific clinical applications.

## Introduction

Mesenchymal Stromal Cells (MSCs) are widely investigated as regenerative therapy in human and veterinary medicine, based on their capacity for tissue remodelling and immunomodulation. However, translation of MSCs into clinical practice has been problematic due to heterogeneity of MSC populations^1^ that is influenced, among others, by cell donor characteristics.^2,3^ Advancing knowledge of the donor-related effect could improve standardisation of MSC therapy through purposeful selection of cell sources, and identify cases when an allogenic approach may be beneficial.

Ageing leads to a decline in function of all cell types, making donor age one of the main considerations in studying MSC variability. MSC therapy is often used in an autologous manner to address potential ethical and safety concerns related to the use of allogenic cell sources.^2,3^ Age is also a risk factor for diseases investigated as targets for MSC therapy such as osteoarthritis,^4^ tendinopathy,^5^ neurodegeneration^6^ or macular degeneration.^7^ Consequently, MSCs applied in a clinical setting are likely to be isolated from older patients, highlighting the importance of understanding the effect of age on MSC potency.

Donor age and sex affect multiple MSC functions including proliferation and differentiation,^8–15^ immunomodulation,^16^ gene expression,^14,17–19^ DNA methylation^20,21^ and energy metabolism.^20^ However, the effect of donor characteristics on protein secretion in MSCs is less well explored. Secreted factors are the main mechanism mediating MSC action and have been investigated as a potential cell-free regenerative therapy.^2,22^ One of the key features of cellular ageing is development of the senescence-associated secretory phenotype (SASP) that can induce age-related pathologies in surrounding tissue. SASP markers identified following induction of senescence in fibroblasts and epithelial cells^23^ were used to test the effect of ageing on MSC protein secretion.^24–28^ However, as SASP components vary across cell types,^29^ comprehensive proteomic analysis is required to characterise the age-related changes in MSC protein secretion.

Untargeted proteomics was previously used to investigate differences in composition of conditioned media (CM) from MSCs isolated from various tissue sources^30–33^ and from different passages of a long-term cell culture, serving as an *in vitro* model of ageing. ^33–37^ *In vitro* ageing does not reflect the complexity of chronological ageing, and these two processes were previously shown to have a different impact on the function of rat adipose-derived MSCs.^38^ Therefore, MSC data from donors of different ages is needed to fully assess the relationship between MSC ageing and secreted proteins.

This study aimed to test the effect of donor sex, age and tissue source on the proteins secreted by equine MSCs. We combined targeted and untargeted proteomics to investigate association between donor characteristics and the abundance of proteins in MSC CM. The results demonstrate that donor age and sex are not likely to drive global changes in MSC CM proteome, however, donor age affects the abundance of individual proteins that may be relevant to tissue repair.

## Materials and Methods

### Cell collection and culture

The owners of all the horses included in the study have been made aware that tissues may be retained for research purposes and provided informed consent for their use in research. Bone marrow-derived MSCs (BMSCs) were obtained for the purpose of autologous cell therapy in privately owned horses and provided by Biobest Laboratories Ltd. Surplus samples of BMSCs from treatment were cryopreserved and stored for research use. Equine adipose-derived MSCs (ASCs) were isolated from retroperitoneal adipose tissue obtained as a waste tissue from colic (abdominal) surgery in privately owned horses, where it was necessary to remove small quantity to aid closure of laparotomy incision. Tissue collection for research was approved by the Committee on Research Ethics, University of Liverpool School of Veterinary Science (RETH000689).

All tissue culture reagents were sourced from Gibco. Adipose tissue was placed in cold phosphate buffer saline (PBS) with 1% penicillin/streptomycin and processed within 12h as previously described.^39^ ASCs were isolated following tissue digestions with collagenase type I (Supplementary File 1) and cultured in T75 flask in an expansion culture medium, consisting of low-glucose Dulbecco’s modified Eagle’s medium (DMEM), supplemented with 30% foetal bovine serum (FBS), 100 U/ml penicillin/streptomycin and 2.5 µg/ml of amphotericin B. After reaching 70% confluence ASCs were passaged, cryopreserved and stored in liquid nitrogen. In all experiments, cells were used at passage 2 or 3, unless stated otherwise. In all cell culture experiments medium was changed twice a week unless stated otherwise. The cells were cultured at 37°C in humidified atmosphere with 5% CO_2_ and 5% O_2_ (hypoxic conditions).

### Tri-lineage differentiation assay

All MSC samples were assessed for their ability to differentiate into osteoblasts, adipocytes and chondrocytes following induction with commercial differentiation media (StemPro, Gibco) previously applied in equine MSCs.^40^ Adipogenic and osteogenic differentiation were undertaken in a monolayer culture for 14 days while chondrogenic differentiation was tested in pellet culture^41^ for 21 days. Cells cultured in expansion medium were used as a negative control and results assessed through specific stains (Sigma-Aldrich); Alizarin red for osteogenic, Oil Red O for adipogenic and Alcian Blue for chondrogenic differentiation. Details of differentiation assays are included in Supplementary File 1.

To compare the size of cell pellets, microscopic images of pellet cultures were taken before processing for histology, and analysed with ImageJ.^42^ The area of the cell pellet and the field of view (FOV) were measured and pellet size reported as a fraction of FOV. The difference between the size of chondrogenic and control pellet for each tissue type was analysed with a paired t-test and means considered significantly different when related p-value was <0.05.

### Expression of MSC markers

RNA from each MSC sample was extracted with TRIzol reagent (Invitrogen) and RNeasy Mini Kit (Qiagen) following manufacturers protocols. Reverse transcription was performed using 1µg RNA and 0.5µg random primers, 200U of M-MLV reverse transcriptase, 25U of RNAsin ribonuclease inhibitor and 500µM dNTP deoxynucleotide mix (all reagents from Promega). Gene expression of CD90, CD29, CD44, CD105, CD34, CD45, CD79α and glyceraldehyde-3-phosphate dehydrogenase (GAPDH) was measured by RT-qPCR using Takyon SYBR Master Mix (Eurogentec), 100nM of specific primers, using previously published sequences^43^ (Table 1), and Roche Lightcycler 480 (Roche). Expression of MSC marker genes relative to GAPDH was calculated using the comparative Ct method.^44^ Sample without the RNA template added in reverse transcription served as a negative control.

**Table 1.**
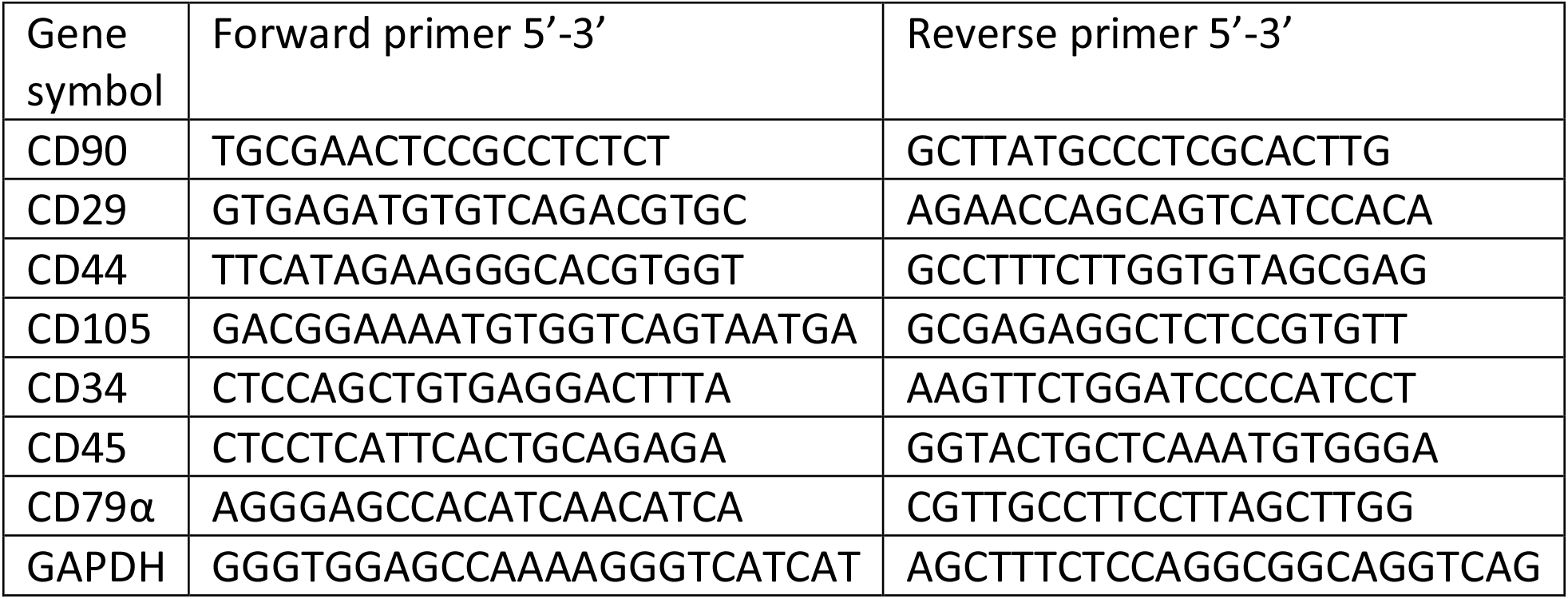
Primer sequences used for equine MSC characterisation.

### Stable Isotope Dynamic Labelling of Secretomes (SIDLS)

To distinguish proteins secreted by MSCs from the structural proteins released into culture media through cell damage, we used dynamic labelling with stable isotopes.^45^ This method used the incorporation of stable isotope-labelled amino acids into newly-made proteins released from cells over time, to infer their rate of synthesis and subsequent secretion. For this experiment, one sample of BMSCs at passage four was seeded in four T25 flasks at 5,000 cells/cm^2^. At 80-90% confluence, expansion media was removed, cells washed 3 times with PBS and 5ml of serum-free DMEM deficient in lysine and arginine (Gibco) with (13C6)-labelled L-lysine (CK Isotopes Ltd, UK) and (13C6/15N4)-labelled arginine (CK Isotopes Ltd, UK) was added to each flask. Conditioned media (CM) was collected after 1h, 2h, 6h or 24h incubation (one flask per time point), centrifuged at 800 x *g* for 7 min to remove cell debris, flash frozen and stored at −80◦C prior to mass spectrometry analysis.

### Label-free conditioned media (CM) collection

Label-free conditioned media were obtained from all MSC samples to investigate the association between protein abundance and donor age, sex and tissue source. Each MSC sample was seeded into two T25 flask at 5,000 cells/cm^2^. CM from one of the replicates was used for mass spectrometry analysis while the other for validation of the results with western blotting and an enzymatic activity assay. At 80-90% confluence, expansion media was removed, cells washed three times in PBS and 5ml of serum-free expansion media added. After 24h incubation media samples were collected, centrifuged at 800 x *g* for 7 min to remove cell debris, flash frozen and stored at −80◦C.

### Protein concentration with StrataClean

Isotope labelled and label-free media were analysed in two separate mass spectrometry experiments following the protocol described below, unless stated otherwise. Protein concentration in CM samples was measured with Pierce 660nm assay (Thermo Scientific) following the manufacturer’s protocol. Protein was concentrated using the StrataClean resin (Agilent Technologies).^45^ CM volume for StrataClean binding was adjusted to obtain 100µg of total protein. In samples with lower protein concentrations, the whole volume of 5ml CM was used. Each CM sample was mixed with 12 µl of StrataClean resin, vortexed for 1 min, then mixed by inverting for 10 min. Samples were centrifuged at 295 x *g* for 1 min and protein-depleted supernatants discarded. StrataClean resin was washed with LC-MS grade water (Thermo Scientific) and centrifuged at 295 x *g* for 1 min, twice, followed by on-bead protein digestion.^45^

### Protein digestion for LC-MS/MS

The on-bead protein digestion with trypsin and lysyl endopeptidase (LysC) was performed as described previously. ^45^ Details of experimental method are included in Supplementary File 1.

### LC-MS/MS spectral acquisition

Mass spectrometry analysis of the secretome samples was undertaken in the Centre for Proteome Research at the University of Liverpool. Protein digest samples were analysed using an Ultimate 3000 RSLC™ nano-system (Thermo Scientific) coupled to a Q Exactive HF Quadrupole-Orbitrap™ mass spectrometer (Thermo Scientific) and a 2-hour gradient. Details of analytical method are included in Supplementary File 1.

### Protein identification and quantification

Mass spectrometry data from two experiments were analysed separately using MaxQuant software (v1.6.17.0 [SIDLS] and v2.0.3.0 [label-free], Max Planck Institute of Biochemistry).^46–48^ Proteins were identified using UniProt reference proteome for *E. caballus* (9796, accessed 11.05.2021 and 10.05.2022, 20,861 entries) and Andromeda search engine.^48^ MaxQuant default search parameters were used (Supplementary File 1). Peptide and protein false discovery rate (FDR) were set to 1% for reporting of peptide spectrum matches (PSM) and protein groups.

Additionally, in analysis of SIDLS data, labels Lys8 and Arg10 were selected to enable identification of labelled peptides. In analysis of label-free data, label-free quantification (LFQ), a MaxQuant intensity determination and normalization algorithm, was enabled.^46^

### Kinetic analysis of secreted proteins identified with SIDLS

Data from SIDLS experiment was analysed following the previously published method^45^ using the R (v4.1.0) programming language environment unless stated otherwise. MaxQuant ‘evidence.txt’ search results file was used as input data. First, reverse and contaminant peptides were removed. For each identified peptide, at each time point, the relative isotope abundance (RIA) was calculated by dividing abundance of the isotope-labelled (heavy) peptide by summed abundance of labelled and non-labelled (light) peptide (RIA = H/[H+L]). To model protein labelling over time (RIA trajectory), we used only peptides identified and quantified in at least three of the four time points. To obtain protein-level kinetic data, we grouped RIA data for peptides assigned to the same protein (‘Leading razor protein’). For each peptide subset, changes of RIA over time were modelled as a first order rate process and fitted via non-linear least squares as described previously.^45^ Next, we calculated the difference in the total mean abundance of peptides (H + L), for each protein, between 24h and 6h time point (P) and used this to calculate protein flux from the cell to extracellular space, by multiplying this with the first-order rate constant (*k*) at which each protein acquired isotopic label. To differentiate secreted from intracellular proteins, we plotted the rate of label incorporation (*k*) against log10 flux for each protein and empirically selected a cut-off value of *k* to separate and classify proteins with distinct kinetic properties.

### Validation of kinetic protein classification

To validate protein classification based on *k* and their flux through the protein pool, we cross-annotated the protein list with UniProt Gene Ontology (GO) terms for cellular component (CC) and undertook GO enrichment analysis of the secreted and intracellular protein subsets using the R package clusterProfiler (function enrichGO(), v.4.2.2).^49,50^ The list of all proteins identified across all four time points (271) was used as a background for over-representation analyses. Gene symbols were used as an input and searched against human gene annotation database (‘org.Hs.eg.db’) due to lack of species-specific database supported by R.

### Clustering analysis of label-free mass spectrometry data

For label-free dataset analysis, MaxQuant output file ‘proteinGroups.txt’ was used. Reverse and contaminant hits were removed and only proteins (protein groups) with two unique peptides retained for statistical analysis.

Before commencement of the clustering analysis, we first investigated the quality of the data by visualising the missing protein intensity values. In order to allow for a reliable analysis using a more complete dataset, we removed proteins that showed more than 30% missing values across all samples. To further investigate the impact of missing values on our analysis, we compared the effects of using different imputation methods on our clustering methods; i) no imputation, ii) sample-based imputation and, iii) protein-based imputation. A detailed explanation of these methods is included in the Supplementary File 1.

We started clustering analysis with an unsupervised overview of the data using principal component analysis (PCA) to compare the effects of tissue type and sex. We then analysed each tissue separately, and clustered the samples based on age. As the ages of the samples were variable and few samples shared ages with other samples, it was decided to group the samples in age range bins of five-year intervals. This allowed identification of clustering patterns associated with age as each group consisted of at least three samples.

Next, we used time course clustering to purposefully identify co-expressed genes across different ages. We used two similar methods; dynamic time clustering (using the R package DTWclust v1.23-1)^51^ and fuzzy time course clustering (Mfuzz v2.56.0).^52^ We first took the mean protein abundance for duplicated samples (i.e. samples with the same tissue type, sex and age). Average values per sample were then standardised at a protein level and the samples ordered by age. For DTWclust, eight clusters were used and the proteins assigned to each cluster identified. Mfuzz was used in a similar way, this time specifying 16 clusters. From the time course clustering approaches, it was hoped we aimed to identify trends in protein expression over time (i.e. age) for each of the assigned clusters.

Finally, an unsupervised clustering method, similar to PCA, was used to identify what drove the variation in protein abundance within tissues. This was completed using with non-negative matrix factorisation (NMF), using the R package NMF (v0.25).^53^ In order to choose the cluster number, we used non-smooth NMF testing between two and six clusters, in order to evaluate the stability of the cluster allocations. From this, we then chose to complete the NMF using four clusters, as this showed the most stable results. We then visualised the association of clusters with each of the known variables (donor age, sex, tissue source).

### Generalised Linear Models of protein abundance

To investigate associations between the abundance of secreted proteins and donor age, sex and tissue type, a generalised linear model (GLM) was applied to each protein previously identified as secreted using SIDLS, and also identified in the label-free dataset obtained from 30 independent MSC samples. The observed protein abundance was modelled as a Gamma distribution to reflect the positive-only and skewed data. A log-link was used, and the analysis was performed using R v.4.1.1. The effect of predictors was considered significant when t-statistics correspond to p<0.05, following adjustment for multiple testing with Benjamini-Hochberg method (q<0.05). Samples with missing data were excluded from analysis on an individual protein basis and information regarding the sample size used in each model reported. For proteins where a categorical predictor (sex, tissue type) was considered to have a significant effect on abundance, the log2 fold change in abundance between conditions was calculated (BMSC relative to ASC, male relative to female).

### Lysyl oxidase activity assay

The activity of lysyl oxidase (LOX) in CM samples (n = 27) was measured using fluorometric commercial assay (Lysyl Oxidase Activity Assay Kit, Abcam) following the manufacturer’s protocol. The relationship between LOX activity and donor age was evaluated using generalised linear mixed-effects models with incubation time, age and a time-by-age interaction term as fixed effects, individual donor as a random effect and normalised fluorescence as the dependent variable (Supplementary File 1).

### Western Blot analysis of collagen triple helix repeat-containing protein 1 (CTHRC1)

CM samples with the highest and lowest CTHRC1 abundance, according to the LC-MS/MS analyses, from each tissue source, were selected for western blotting (n = 6). Samples were concentrated using StrataClean resin as described previously, ^54^ subjected to Sodium dodecyl-sulfate polyacrylamide gel electrophoresis and stained with rabbit anti-human CTHRC1 antibody (Abcam, ab85739). Fibronectin detected using an anti-human antibody (Sigma-Aldrich, F3648) was used as a loading control.^55^ To quantify the CTHRC1 abundance, the relative density of CTHRC1 band (ratio of CTHRC1 to FBN) was evaluated using ImageJ.^42^ Details of the experimental methods can be found in Supplementary File 1.

### ELISA analysis of monocyte chemoattractant protein 1 (MCP-1)

There was insufficient information regarding the abundance of a dominant marker of SASP, MCP-1, in the mass spectrometry dataset (>30% missing values). Therefore, we have used a commercial sandwich enzyme-linked immunosorbent assay (Horse MCP1 ELISA Kit, Abcam) to measure concentration of MCP-1 in MSC CM (n = 24) following manufacturer’ protocol. The relationship between donor profile and tissue source and MCP-1 concentration was tested using GLMs as described for the mass spectrometry data (Supplementary File 1).

## Results

### Characterisation of equine Mesenchymal Stromal Cells

This study used equine MSCs isolated from bone marrow (n = 14) and adipose tissue (n = 16) collected from horses aged from 1.5 to 24 years (mean 11.7 years), 14 males and 16 females (Figure 1A). MSCs from both tissue sources showed positive relative expression (RE) of genes encoding MSC surface marker proteins CD90 (mean = 0.79), CD29 (mean = 0.17), CD44 (mean = 0.17) and CD105 (mean = 0.05), and low expression of CD34, CD45 and CD79α (mean < 0.001), indicating that the cell populations used did not include hematopoietic stem cells or B cells (Figure 1B). Staining of MSC cultures with Alizarin Red following osteogenic induction showed the presence of calcium deposits and staining with Oil Red O showed the presence of lipid droplets in adipogenic cultures (Figure 1C). The mean size of MSC pellet cultures maintained in chondrogenic media was 2.5- (ASC) and 3.5- (BMSC) times higher than control pellets (p < 0.05, in both tissue types, Figure 1D). Paraffin sections of chondrogenic MSC pellets showed positive staining for Alcian Blue, indicating presence of glycosaminoglycans (Figure 1C).

**Figure 1.**
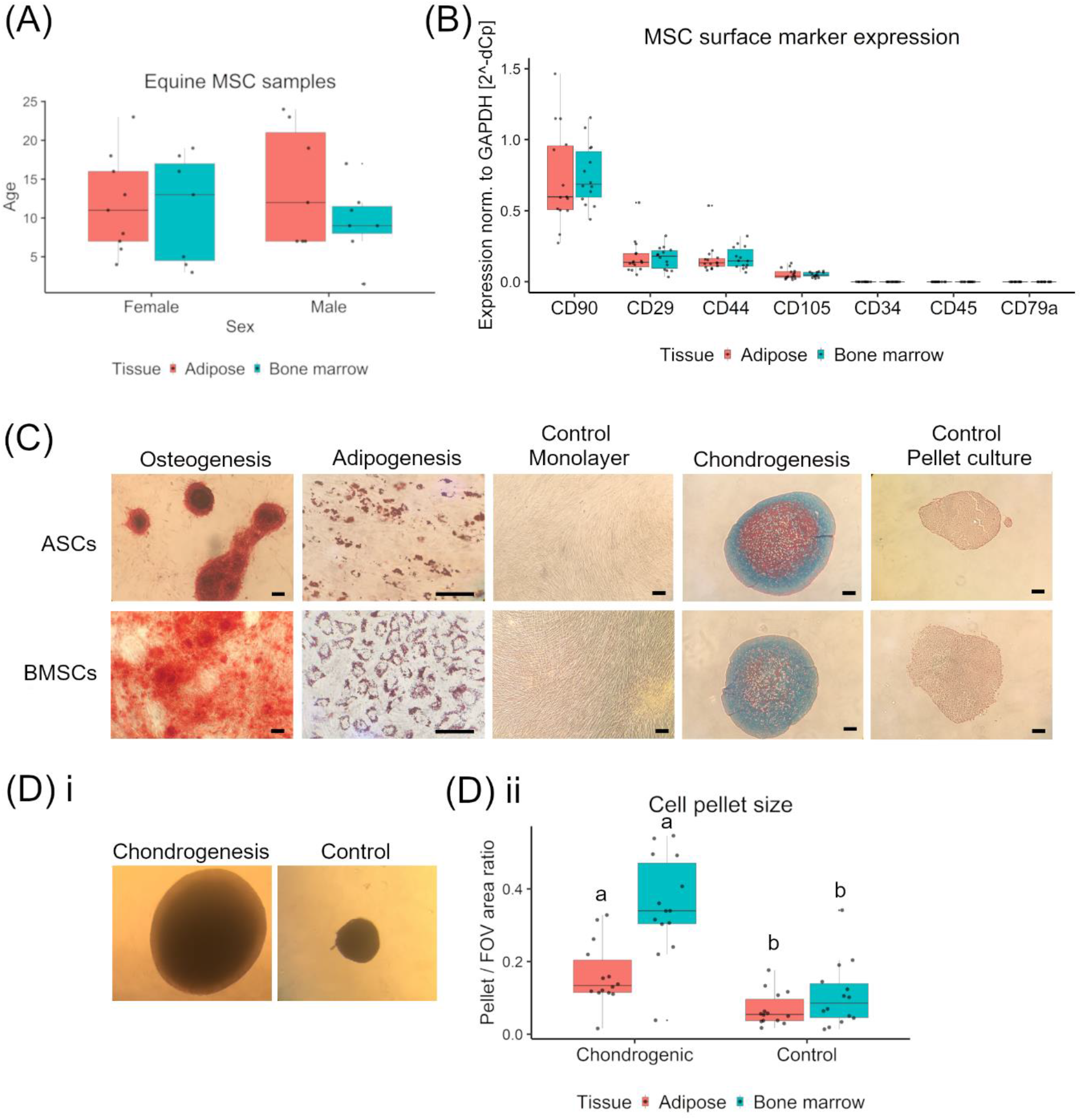
Characterisation of equine mesenchymal stromal cells (MSC). A, Distribution of MSC samples (N = 30) with regard to donor age, sex and tissue type used for cell isolation. B, Expression of MSC surface marker genes relative to glyceraldehyde-3-phosphate dehydrogenase (GAPDH), measured with qRT-PCR. C, Microscopic images of adipose- and bone marrow-derived MSCs grown as monolayers in osteogenic induction media and stained with Alizarin Red; adipogenic induction media and stained with Oil Red O; grown in pellet culture in chondrogenic induction media, paraffin sectioned and stained with Alcian Blue. Control cultures were grown in MSC expansion media. Representative images from six biological replicates. Bar; 100 μm. D, Representative microscopic images (n = 1) used for MSC pellet size measurements (i) and graph showing relative size of pellets grown in chondrogenic and control media (ii). Different letters indicate statistically significant (P < 0.05) differences between chondrogenic and control group in samples from the same tissue source, determined with paired t-test; (ASC, n = 14; BMSC, n = 14). In all box plots horizontal lines represent median, boxes interquartile range, whiskers range and dots biological replicates.

### Stable isotope labelling identified secreted proteins in MSC CM based on their kinetic behaviour

Out of 163 proteins with peptides identified in CM samples in at least three of four time points, the rate of synthesis and secretion (*k*), and flux, for 106 were characterised in detail (Figure 2A, Supplementary File 1). Based on an empirically selected cut-off value of *k* = 0.01, 70 proteins were classified as secreted and 36 as intracellular (Figure 2B). Differences in dynamic behaviour between secreted and intracellular proteins underlying this classification are illustrated in Figure 2C.

**Figure 2.**
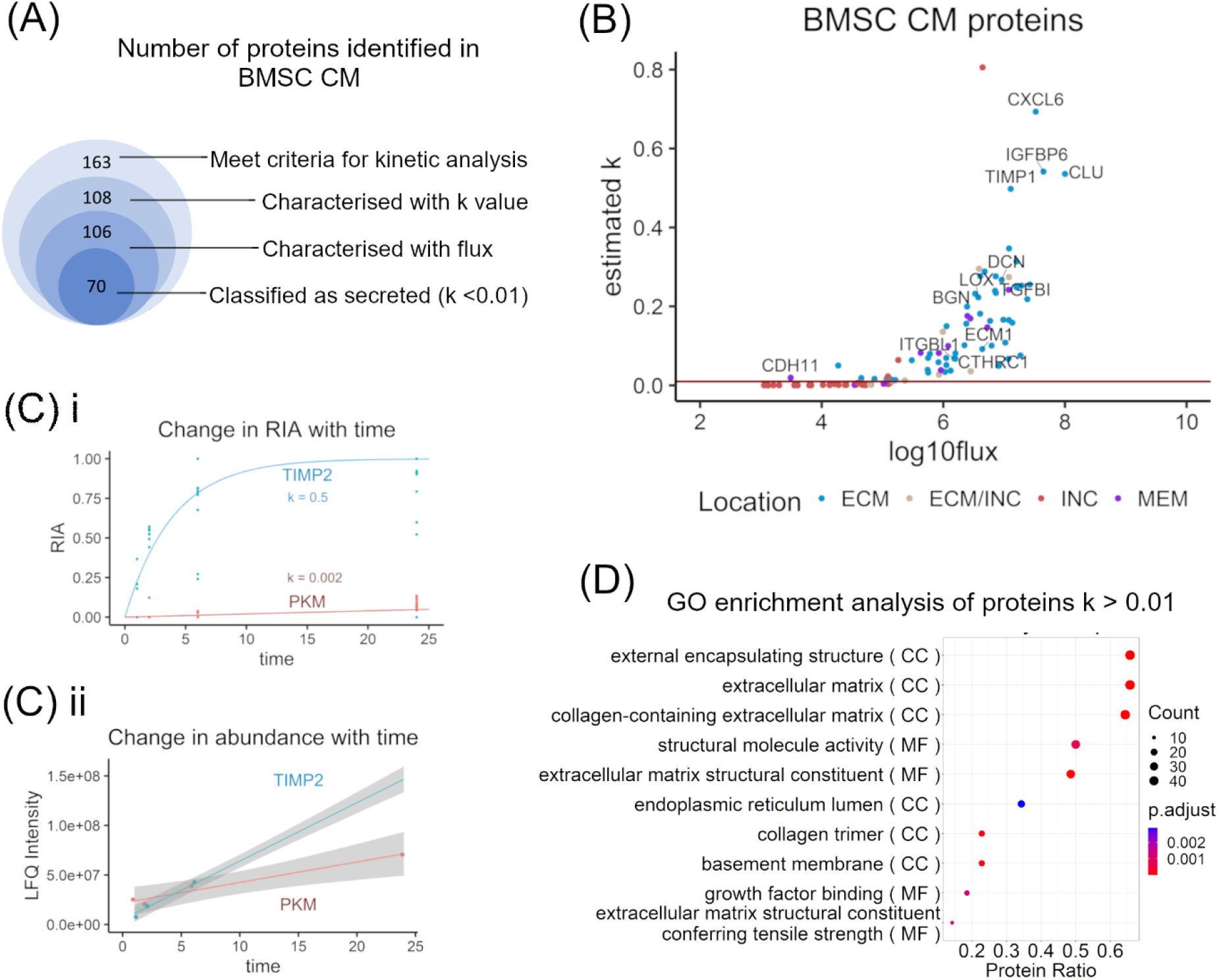
Stable Isotope Dynamic Labelling of Secretomes (SIDLS) allowed identification of classically secreted proteins in conditioned medium (CM) of equine Mesenchymal Stromal Cell (MSCs). A, Stacked Venn diagram showing number of proteins at consecutive stages of mass spectrometry data analysis. The innermost circle shows the number of proteins characterised as secreted, based on their kinetic properties. B, Scatterplot showing 106 proteins identified in MSC CM and characterised with kinetic parameters: first-order rate constant (*k*), describing change of Relative Isotope Abundance in time, and flux, describing the rate at which the protein flows from the intracellular to the extracellular space. Red line represents empirically selected threshold (*k* = 0.01) discriminating proteins with distinct kinetic behaviour. Colour-coding was based on Gene Ontology (GO) cellular component (CC) terms. Labels designate proteins with the highest secretion rate and those that were influenced by donor characteristics in this study. C, Dynamic behaviour of representative secreted (Metalloproteinase inhibitor 1, TIMP1) and intracellular (pyruvate kinase, PKM) proteins. The Relative Isotope Abundance (RIA) describes the ratio of heavy-labelled peptide abundance to the sum of labelled and non-labelled peptide abundances. TIMP1 exists in a small intracellular pool, acquires heavy-labelled amino acid quickly during nascent *de novo* protein synthesis, while the intracellular pool of PKM is large and acquires heavy labels over much longer time-periods. Dots represent RIA values of single peptides and the lines the fitted curves for a first order equation (i). Total abundance of secreted proteins in CM increases in time faster than intracellular proteins derived from cell leakage/death. Dots represent mean protein abundance with regression lines and 95% confidence interval (shaded area) (ii). D, Dot plot of GO term enrichment analysis of 70 proteins classified as secreted based on kinetic properties. The size of the dots indicates the number of proteins mapped to that term. The x-axis shows number of proteins that map to the term divided by the total number of secreted proteins. The dots are colour-coded by adjusted p-values (BH method).

Cross-annotation with GO cellular component terms showed that, out of 70 classified as secreted, 58 were confirmed as extracellular proteins, five of intracellular origin, one a membrane protein and six were assigned to multiple (intra- and extracellular) locations. All proteins with *k* < 0.01 were assigned to the intracellular pool or multiple locations (Figure 2B). As expected, GO enrichment analysis of proteins with *k* > 0.01 resulted in top cellular component and molecular function terms associated predominantly with extracellular matrix (ECM) structure (Figure 2D). The full list of 106 proteins and their kinetic parameters is available in Supplementary File 2.

### Tissue origin is the main factor affecting protein composition of equine MSC CM

Label-free quantification analysis of equine MSC CM samples from 30 donors identified 1164 proteins based on the presence of minimum two unique peptides. Following filtering for proteins where quantification values were missing in over 30% of samples, 716 proteins were used for clustering analysis using non-imputed, sample-imputed and protein-imputed datasets, which showed that imputation did not affect the sample clustering patterns (Supplementary File 1, Supplementary Figure 1). Therefore, we decided to present results of the analysis using only the non-imputed variables (314 proteins).

PCA showed that tissue origin of MSCs is the main variable driving clustering of CM samples, accounting for up to 52% of variance in protein abundance (Figure 3A). There was no clear association between donor sex or age and sample clusters, including after removing the tissue factor (Figure 3A, Supplementary Figure 2A). The unsupervised clustering using NMF method resulted in four distinct sample clusters. The number of clusters was assigned based on the NMF clustering rank validation (Supplementary Figure 2B). One of the clusters (cluster 4) included 13 out of 14 BMSC CM samples while ASC CM samples were grouped into three sub-clusters (Figure 3B), supporting the finding that tissue type is driving similarity in protein abundance across MSC CM samples. NMF clusters did not appear to be associated with sex or age of the donor (Figure 3Ci). Top proteins driving similarity between samples within each cluster included mainly extracellular (secreted) proteins, apart from the cluster 2, that was characterised by cytosolic and membrane proteins (Figure 3Cii). The full list of protein ranks is available in Supplementary File 3.

**Figure 3.**
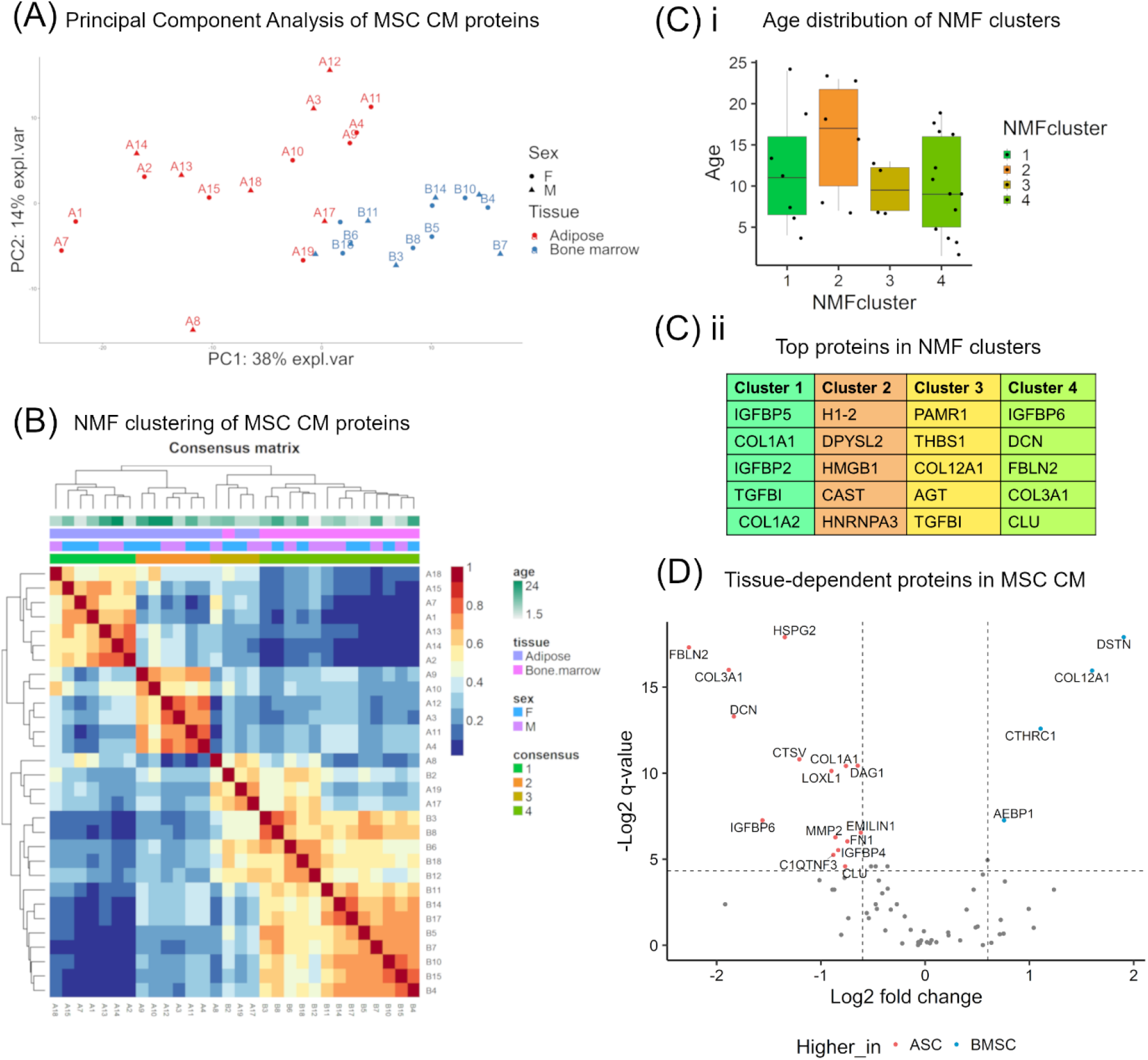
Tissue source is the main factor associated with differences in the abundance of secreted proteins in equine mesenchymal stromal cell (MSC) conditioned media (CM). A, Principal Component Analysis (PCA) biplot of label-free mass spectrometry data obtained from MSC CM samples (n = 30) showing the first two principal components. Each dot represents a biological replicate with colour denoting the tissue type used for MSC isolation and shape donor sex. B, Consensus matrix from non-negative matrix factorisation (NMF) clustering of MSC CM samples based on the normalised abundance values (LFQ intensity) of 314 proteins identified in all biological replicates with mass spectrometry. Four distinct clusters were identified (colour coded as ‘consensus’) and mapped against MSC source characteristics (colour coded age, tissue and sex). C, i) boxplots showing distribution of donor age in samples allocated to four NMF clusters; ii) Gene names of the top proteins identified in each NMF cluster (colour code consistent with consensus matrix). Abundance of those proteins was the main source of similarity between MSC CM samples allocated to each cluster. D, Volcano plot showing 69 proteins classified as secreted by SIDLS that were further identified with label-free mass spectrometry in MSC CM samples (n = 30). The x-axis shows log2 fold change in mean abundance of each protein (dots) in bone marrow-relative to adipose-derived MSC CM. The y-axis shows the -log2 q-value (FDR adjusted p-value, BH method) associated with the effect of tissue source on protein abundance as determined with a generalised linear model (GLM). Proteins with log2 fold change > 0.6 or < −0.6 and p-value < 0.05 were colour-coded and labelled with gene names.

Out of 70 proteins classified as secreted by SIDLS, 69 were identified in label-free dataset and were progressed to the association study using GLMs described above. The null hypothesis of no effect of tissue origin could be rejected at the level of 0.05 for 23 proteins (Supplementary File 4), and for 19 of these proteins the log2 fold change in mean protein abundance between tissue groups was larger than 1.5 (log2 fold change > 0.6 or < −0.6, Figure 3B). Following adjustment for multiple comparisons (q-values) and log2 fold change, when applying a significance threshold of 0.05, 15 proteins were considered upregulated in ASC CM while four were upregulated in BMSC CM (Figure 3B). All of the proteins previously identified as most characteristic of BMSC CM cluster in NMF analysis were significantly downregulated in BMSC CM according to GLM and fold change analysis (Figure 3B).

### Abundance of proteins related to ECM remodelling declines with donor age in equine MSC CM

Time course clustering using the MFuzz approach, with age as a time factor, identified 16 clusters of co-expressed proteins. (Figure 4A, Supplementary File 5). However, protein abundance trajectories in those clusters did not show correspondence with donor age. Similarly, dynamic time warping (DTW) with 8 clusters assigned did not identify a clear trend in protein cluster abundance related to donor age (Supplementary Figure 2C, Supplementary File 6). Varying the clusters for Mfuzz and DTWclust did not affect the main observation of lack of correspondence between clusters and age trajectory.

**Figure 4.**
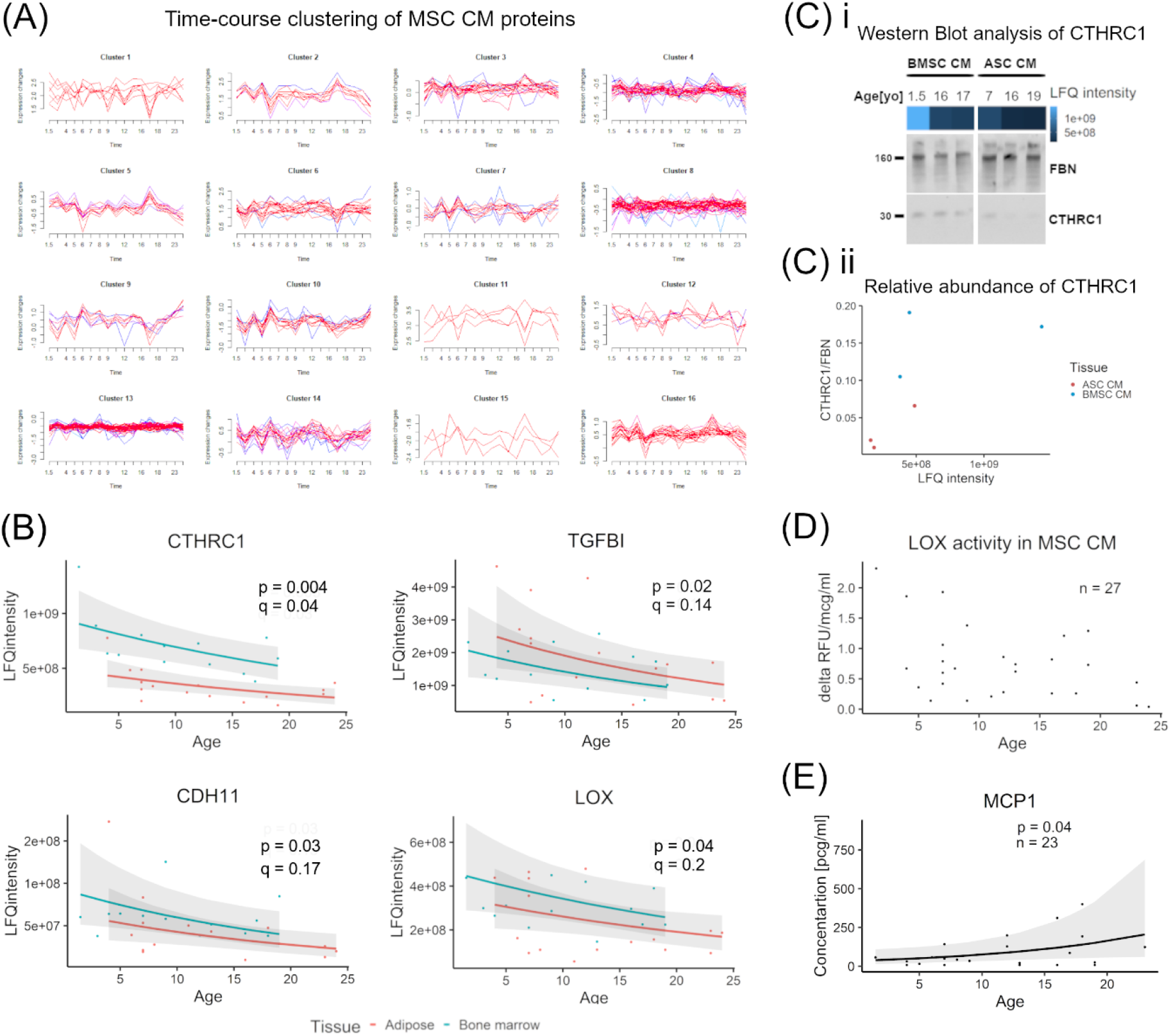
Donor age affects the abundance of selected proteins identified in equine mesenchymal stromal cell (MSC) conditioned media (CM). A, Cluster members identified with fuzzy time-course clustering. Graphs show clusters of proteins with similar abundance patterns and change in their abundance value with donor age. B, Scatterplots showing change in CM protein abundance, measured by quantitative mass spectrometry, with MSC donor age. Dots represent biological replicates, colours tissue type used for MSC isolation, lines the mean protein abundance predicted by generalised linear model (GLM) and shades 95% confidence interval (CI). Proteins with the strongest relationship between age and abundance (p-value <0.05, q-value <0.2) are shown: collagen triple helix repeat-containing protein 1 (CTHRC1), lysyl oxidase (LOX), transforming growth factor beta induced (TGFBI), cadherin 11 (CDH11). C, Western blot analysis of the abundance of CTHRC1 in CM from bone-marrow-derived (BMSC) and adipose-derived (ASC) equine MSCs, isolated from donors of different age (n = 6). Heatmap bar shows the abundance of CTHRC1 (LFQ intensity) measured in CM by mass spectrometry. Fibronectin (FBN) was included as loading control (i). Density of CTHRC1 protein bands relative to FBN were measured with ImageJ and plotted against LFQ intensity of CTHRC1 in CM from the same MSC samples (ii). D, Scatterplot showing enzymatic activity of LOX in MSC CM (n = 27), measured with a commercial fluorescence assay. The y-axis shows the difference in fluorescence intensity (RFU – relative fluorescence units), normalised to total protein concentration, between 30 min and 10 min incubation time (delta RFU). E, Scatterplot showing change in MCP-1 concentration in MSC CM in relation to donor age. Dots represent biological replicates, lines the mean protein abundance predicted by GLM and shades 95% confidence interval (CI).

Analysis of individual secreted proteins with GLM detected effect of donor age in four proteins (p < 0.05): Collagen triple helix repeat-containing protein 1 (CTHRC1), cadherin 11 (CDH11), lysyl oxidase (LOX) and transforming growth factor-beta-induced protein ig-h3 (TGFBI). However, following correction for multiple comparisons, only in CTHRC1 met the q-value associated with the effect of donor age met the threshold needed to reject the null hypothesis (q = 0.04) (Figure 4B, Supplementary File 4.). The extracellular proteins, CTHRC1, LOX and TGFBI, have been all assigned a high rank in cluster 3 identified by NMF (Supplementary file 3) while CTHRC1 and TGFBI were assigned to the same DTW cluster (Supplementary file 5). This evidence suggests that CTHRC1, TGFBI and LOX may be co-expressed and show the same direction of abundance change (decline) related to cell donor age.

We validated the abundance change in two of these proteins, CTHRC1 and LOX, using orthogonal techniques (western blotting and ELISA, respectively). Western blot analysis of CTHRC1 showed relative abundance corresponded with quantitative values from the LFQ LC-MS/MS proteomics approach (Figure 4C). Analysis of LOX activity in MSC CM showed a significant effect of time-by-age interaction term (b = −0.05, p-value < 0.001), suggesting that the increase of fluorescence over time that reflects LOX enzymatic activity in MSC CM decreased with donor age (Figure 4D). These data demonstrate that donor age is most likely not driving global changes in MSC CM proteome. However, it does negatively affect secretion of several proteins relevant to cell migration, cell-ECM interaction and collagen synthesis.

### Donor age is associated with increased secretion of the SASP marker MCP1 by equine MSCs

The results of ELISA provide evidence of an association between MCP-1 concentration in MSC CM and donor age (b = 0.08, p-value = 0.04, Figure 4E). The results do not show any associations between the donor sex or tissue source and MCP-1 concentration.

### Limited evidence of the effect of donor sex on protein abundance in equine MSC CM

Results of GLM analysis suggested that abundance of three proteins is higher in CM obtained from MSCs derived from male than female donors, however, following FDR adjustment correction for multiple comparisons, none of these effects met the threshold of q < <0.05 (Figure 5, Supplementary File 4). In order to reject null hypotheses related to the effect of donor sex on the abundance of these proteins, the significance threshold would need to be set at q<0.2.

**Figure 5.**
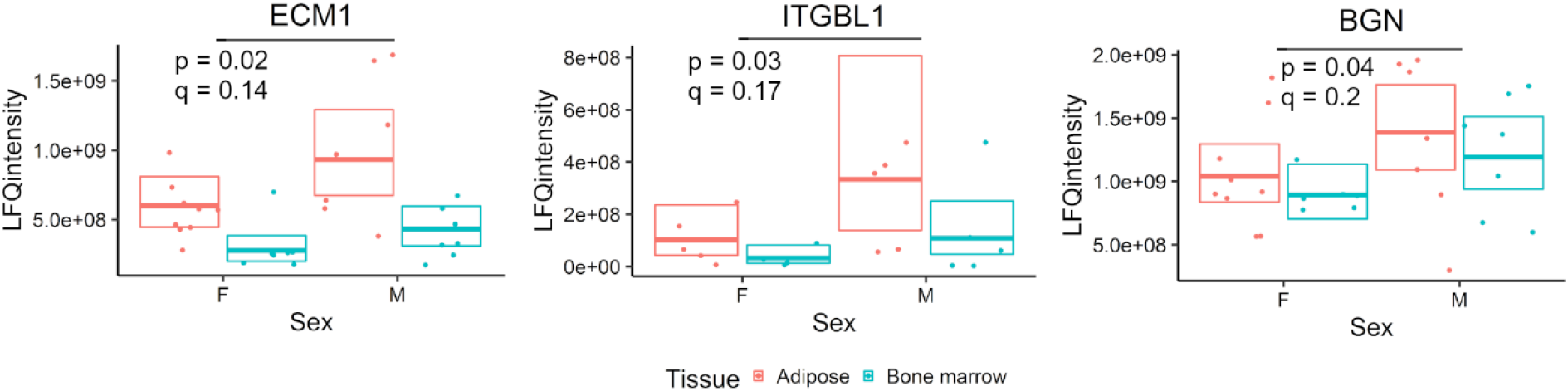
Effect of donor sex on abundance in equine mesenchymal stromal cell (MSC) conditioned media (CM). Dots represent biological replicates, boxes and horizontal lines represent interquartile range and median predicted by generalised linear model (GLM). Colours designate tissue types used for MSC isolation. The proteins where the effect of donor sex on abundance was associated with p-value < 0.05 and q-value < 0.2 are shown: biglycan (BGN), extracellular matrix protein 1 (ECM1) and integrin subunit beta like 1 (ITGBL1).

## Discussion

To our knowledge, this is the first study in any species reporting statistical analysis of a comprehensive protein profile of MSC CM obtained from donors of different age. Mass spectrometry was previously used to analyse the protein composition of ECM synthesised *in vitro* by human BMSCs from young and old donors.^56^ However, this study did not test for systematic differences in protein abundance but focused on proteins identified uniquely in either of the age groups. Other studies testing the effect of ageing on MSC CM all utilised the *in vitro* models of ageing through long-term culture, that are characterised by decreased cell proliferation capacity.^33–37^ Although replicative senescence reflects only one of the aspects of cellular ageing, increasing concentrations of MCP-1 in CM was observed both in MSCs from older horses and late passage human BMSCs.^33–35^ *In vitro* studies showed that the abundance of most CM proteins remains unchanged with progressive cell passaging,^36,37^ however, proteins upregulated in late passage BMSCs had a senescence-inducing effect on early passage cells.^36^ Our results likewise suggests that the abundance of most proteins in equine MSC CM is unaffected by donor age, yet the age-related decrease in CTHRC1, LOX may have functional implications, due to the role of these proteins in regulation of ECM remodelling.

Proteins whose abundance were identified here as age-dependent participate in regulating synthesis of collagen and elastin, the main structural components of ECM. CTHRC1 interacts with the TGF-β/Smad signalling pathway^57^ that controls activation of pro-fibrotic genes. Expression of CTHRC1 has been linked with fibroblasts activation following injury in heart, arterial wall and skin tissue, and with promotion of tissue healing through scar formation^57–59^. LOX is an enzyme mediating post-translational modification (cross-linking) of collagen and elastin that leads to stabilisation of fibre structure, affecting biomechanical properties of the ECM.^60^ Age-related decline in the levels of CTHRC1 in equine MSC CM is in contrast with a recent study in human BMSCs which identified CTHRC1 only in ECM synthesised by cells from old, but not young, donors.^56^ It is possible that our results are specific to equine MSCs; however, it is worth noting that applying data-dependent acquisition mass spectrometry proteomics to the small sample number used in the human study could have resulted in lack of identification of CTHRC1 secreted by young BMSCs. ^61^

The effects seen here of tissue source on equine MSC CM protein content is in line with previous reports,^30–32^ notably, upregulation of CTHRC1 in BMSCs^30^ and clusterin (CLU) and decorin (DCN) in ASCs^31^ which have previously been shown in human MSCs. In addition, our study showed higher variability in ASC secretome composition compared with BMSCs. The ASCs used in this study were isolated from horses undergoing gastrointestinal surgery, while the BMSCs were collected to generate autologous therapy for musculoskeletal disease. Although ASCs were collected only from cases without severe systemic symptoms, the health status of these horses would vary more than that of the BMSC donors. In the absence of any clear associations between known donor characteristics and NMF clusters, this could be considered as one of the potential factors driving variability in the ASC secretome. This finding puts into perspective the previously proposed concept of using fat tissue from laparotomy to isolate ASCs for allogenic therapeutic use.^28,62^

A previous study of sex-related differences in the MSC secretome targeted inflammation-related cytokines measured in human ASC CM^16^. Here, we tested the effect of donor sex on a subset of extracellular signalling proteins characterised as classically secreted in equine MSC CM. Although the evidence from GLM analysis did not meet the threshold needed to reject the null hypothesis, it suggested potential targets for further investigation. In particular, secretion of ITGBL1 could be of relevance to MSC therapeutic potency due to its effect on TGF-β signalling and chondroprotective properties.^63^

Interestingly, six MSC CM proteins that showed kinetic behaviour of a classically secreted protein in the SIDLS experiment were annotated to intracellular and membrane compartments by GO CC terms. However, a review of the literature showed that all of these proteins were previously identified in a secretory form: destrin (DSTN), Tripeptidyl-peptidase 1 (TPP1) and amyloid beta protein (APP) in MSC-derived extracellular vesicles (EVs);^64^ cadherin 11 (CDH11) in tumour-derived EVs,^65^ cathepsin V in secretory lysosomes^66^ and tyrosyl-tRNA synthetase (YARS1) as a soluble cytokine.^67^ This demonstrates the value of dynamic labelling in studying cell secretomes, and especially proteins released by unconventional secretory routes.

To increase the likelihood that our results will be functionally relevant, we have focused on identification and analysis of proteins actively secreted by MSCs. However, conclusions about the functional implications of the present findings are limited to the previous knowledge about the role of those proteins in extracellular niche regulation. Further studies are needed to determine the effect of secretome composition on cell function using relevant *in vitro* models. The results presented here may contribute to our understanding of the variability of the biological effects of MSC through linking protein secretion to fundamental characteristics of a cell donor. Although findings from this study may not translate to humans, it is clear that they raise an awareness for considering the potential clinical implications of MSC donor age, as well as careful consideration of the source of the MSC to be used for treatment.

## Conclusions

Tissue type and donor age contribute to the heterogeneity in protein composition of MSC CM and should be taken into consideration when choosing the source of MSCs used for therapeutic applications.

## Acknowledgments

This project was supported by Horserace Betting Levy Board Equine Post Doctoral Fellowship - VET/2020 -2 EPDF 8. MP was funded by the Horserace Betting Levy Board and Wellcome Trust Clinical Intermediate Fellowship (grant 107471/Z/15/Z).

## Data Availability Statement

Raw data from mass spectrometry proteomic analysis and MaxQuant search output files are available in the ProteomeXchange Consortium via the Proteomics Identification Database (upload in progress).

Code used for mass spectrometry data analysis is available at github.com/CBFLivUni/Turlo_MSC_secretome_proteomics and github.com/aturlo/MSC_secretome_proteomics and results are presented in the supplementary files. Other data that support the findings of this study are presented within the article and its figures.

## Supplementary Files

**Supplementary File 1 (pdf). Details of experimental methods and data analysis**.

**Supplementary File 2 (xmls). List of proteins and their kinetic parameters determined by Stable Isotope Dynamic Labelling of Secretomes (SIDLS) in equine bone marrow-derived mesenchymal stromal cell conditioned media**.

**Supplementary File 3 (xmls). List of clusters and protein ranks assigned based on non-negative matrix factorisation clustering of proteins identified in equine mesenchymal stem cell conditioned media**.

**Supplementary File 4 (xmls). Results of generalised linear model analysis of the relationship between donor age, sex and tissue source and abundance of secreted proteins in equine mesenchymal stromal cell conditioned media**.

**Supplementary File 5 (xmls). List of protein clusters assigned based on dynamic time warping clustering of proteins identified in equine mesenchymal stem cell conditioned media**.

**Supplementary File 6 (xmls). List of protein clusters assigned based on MFuzz clustering of proteins identified in equine mesenchymal stem cell conditioned media**.

## Supplementary Figures

**Supplementary Figure 1.**
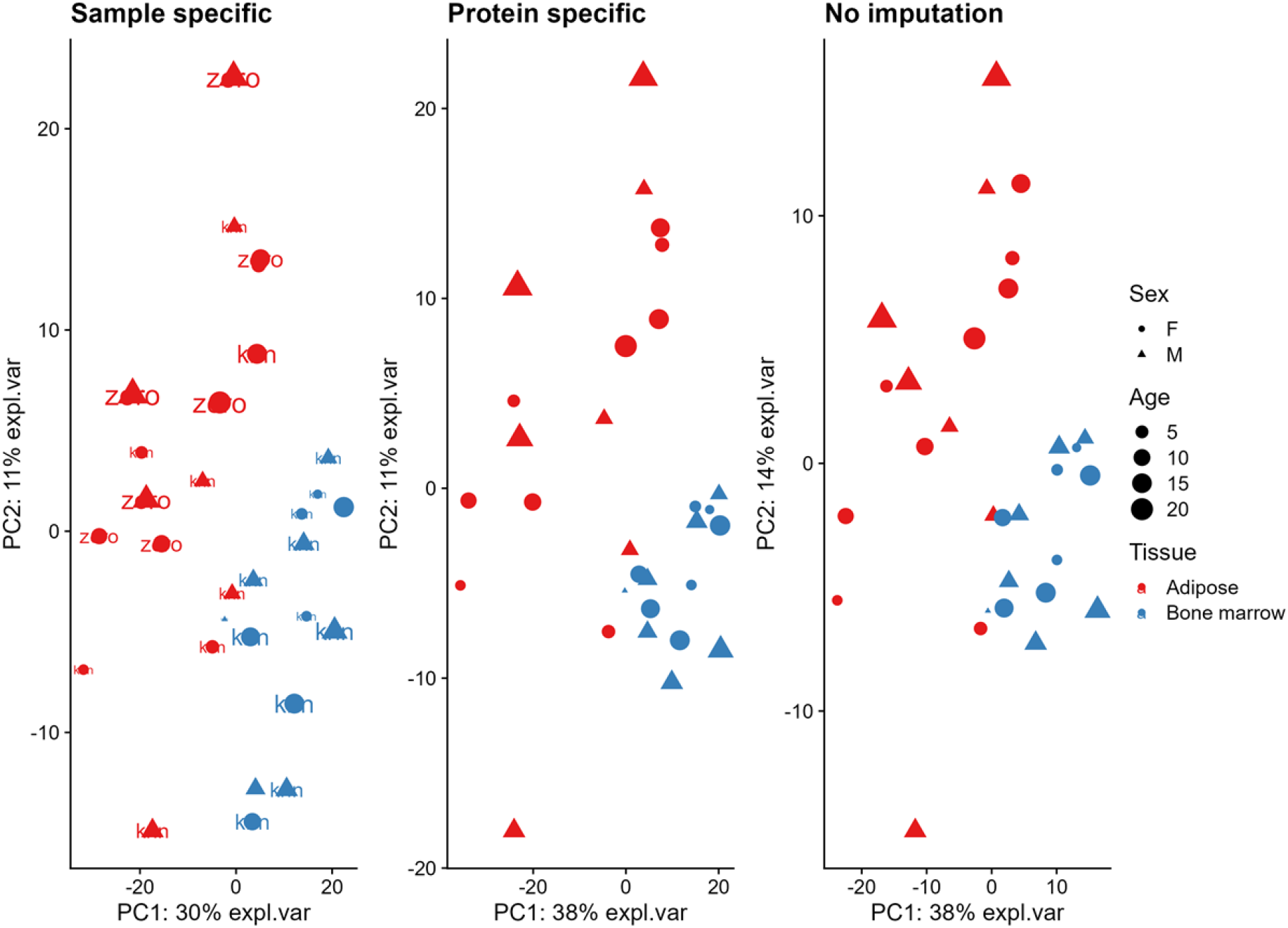
PCA plots of label-free LC-MS dataset resulting from analysis of equine mesenchymal stromal cell conditioned media with different imputation methods and non-imputed.

**Supplementary Figure 2.**
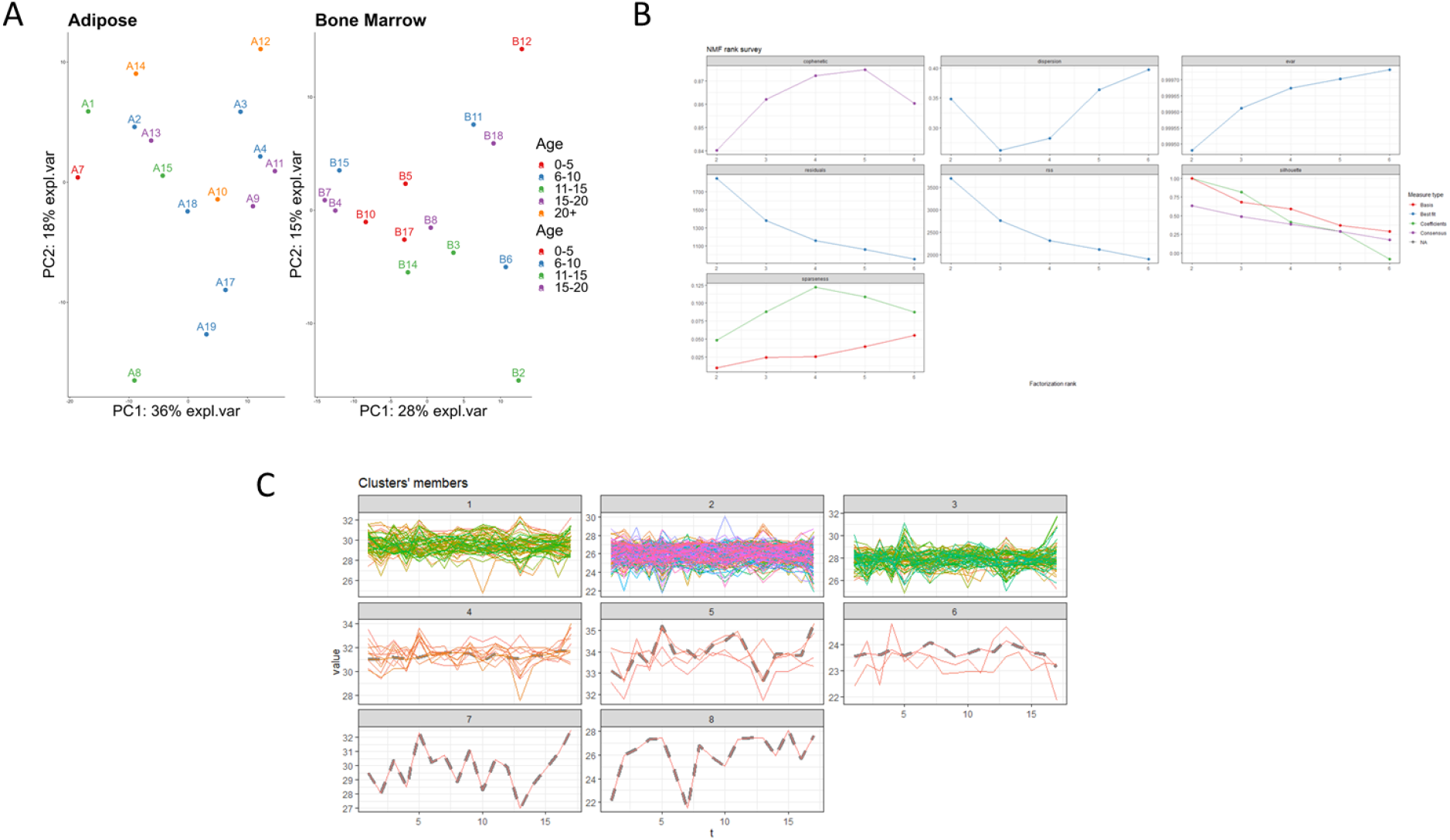
Relationship between equine mesenchymal stromal cell (MSC) donor age and clustering of conditioned media (CM) samples based on their proteomic profile. A, Principal Component Analysis (PCA) of MSC CM by age after removal of the tissue factor. Colour corresponds with MSC donor age range divided in 5-year intervals. B, Non-negative matrix factor (NMF) clustering rank validation to assign cluster number. C, Results of DTW time course clustering of MSC CM proteins.

## References

1. Levy O, Kuai R, Siren EMJ, et al. Shattering barriers toward clinically meaningful MSC therapies. Sci Adv 2020;6(30):eaba6884; doi: 10.1126/sciadv.aba6884.

2. Jovic D, Yu Y, Wang D, et al. A Brief Overview of Global Trends in MSC-Based Cell Therapy. Stem Cell Rev Rep 2022;18(5):1525–1545; doi: 10.1007/s12015-022-10369-1.

3. Wright A, Arthaud-Day ML, Weiss ML. Therapeutic Use of Mesenchymal Stromal Cells: The Need for Inclusive Characterization Guidelines to Accommodate All Tissue Sources and Species. Front Cell Dev Biol 2021;9:632717; doi: 10.3389/fcell.2021.632717.

4. O’Neill TW, McCabe PS, McBeth J. Update on the Epidemiology, Risk Factors and Disease Outcomes of Osteoarthritis. Best Pract Res Clin Rheumatol 2018;32(2):312–326; doi: 10.1016/j.berh.2018.10.007.

5. Guzzoni V, Selistre-De-Araújo HS, Marqueti R de C. Tendon Remodeling in Response to Resistance Training, Anabolic Androgenic Steroids and Aging. Cells 2018;7(12); doi: 10.3390/cells7120251.

6. Hrelia P, Sita G, Ziche M, et al. Common Protective Strategies in Neurodegenerative Disease: Focusing on Risk Factors to Target the Cellular Redox System. Oxid Med Cell Longev 2020;2020:8363245; doi: 10.1155/2020/8363245.

7. Wang Y, Zhong Y, Zhang L, et al. Global Incidence, Progression, and Risk Factors of Age-Related Macular Degeneration and Projection of Disease Statistics in 30 Years: A Modeling Study. Gerontology 2022;68(7):721–735; doi: 10.1159/000518822.

8. Bagge J, MacLeod JN, Berg LC. Cellular Proliferation of Equine Bone Marrow- and Adipose Tissue-Derived Mesenchymal Stem Cells Decline With Increasing Donor Age. Front Vet Sci 2020;7:602403; doi: 10.3389/fvets.2020.602403.

9. Ganguly P, El-Jawhari JJ, Burska AN, et al. The Analysis of in Vivo Aging in Human Bone Marrow Mesenchymal Stromal Cells Using Colony-Forming Unit-Fibroblast Assay and the CD45lowCD271+ Phenotype. Stem Cells Int 2019;2019:5197983; doi: 10.1155/2019/5197983.

10. Li C, Wei G, Gu Q, et al. Donor Age and Cell Passage Affect Osteogenic Ability of Rat Bone Marrow Mesenchymal Stem Cells. Cell Biochem Biophys 2015;72(2):543–9; doi: 10.1007/s12013-014-0500-9.

11. Beane OS, Fonseca VC, Cooper LL, et al. Impact of aging on the regenerative properties of bone marrow-, muscle-, and adipose-derived mesenchymal stem/stromal cells. PLoS One 2014;9(12):e115963; doi: 10.1371/journal.pone.0115963.

12. Zaim M, Karaman S, Cetin G, et al. Donor age and long-term culture affect differentiation and proliferation of human bone marrow mesenchymal stem cells. Ann Hematol 2012;91(8):1175–86; doi: 10.1007/s00277-012-1438-x.

13. Liu M, Lei H, Dong P, et al. Adipose-Derived Mesenchymal Stem Cells from the Elderly Exhibit Decreased Migration and Differentiation Abilities with Senescent Properties. Cell Transplant 2017;26(9):1505–1519; doi: 10.1177/0963689717721221.

14. Alt EU, Senst C, Murthy SN, et al. Aging alters tissue resident mesenchymal stem cell properties. Stem Cell Res 2012;8(2):215–25; doi: 10.1016/j.scr.2011.11.002.

15. Selle M, Koch JD, Ongsiek A, et al. Influence of age on stem cells depends on the sex of the bone marrow donor. J Cell Mol Med 2022;26(5):1594–1605; doi: 10.1111/jcmm.17201.

16. Mckinnirey F, Herbert B, Vesey G, et al. Immune modulation via adipose derived Mesenchymal Stem cells is driven by donor sex in vitro. Sci Rep 2021;11(1):12454; doi: 10.1038/s41598-021-91870-4.

17. Wang B, Liu Z, Chen VP, et al. Transplanting cells from old but not young donors causes physical dysfunction in older recipients. Aging Cell 2020;19(3):e13106; doi: 10.1111/acel.13106.

18. Pandey AC, Semon JA, Kaushal D, et al. MicroRNA profiling reveals age-dependent differential expression of nuclear factor B and mitogen-activated protein kinase in adipose and bone marrow-derived human mesenchymal stem cells. Stem Cell Res Ther 2011;2(6):49; doi: 10.1186/scrt90.

19. Ren S, Xiong H, Chen J, et al. The whole profiling and competing endogenous RNA network analyses of noncoding RNAs in adipose-derived stem cells from diabetic, old, and young patients. Stem Cell Res Ther 2021;12(1):313; doi: 10.1186/s13287-021-02388-5.

20. Toraño EG, Bayón GF, del Real Á, et al. Age-associated hydroxymethylation in human bone-marrow mesenchymal stem cells. J Transl Med 2016;14(1):207; doi: 10.1186/s12967-016-0966-x.

21. Lu GM, Rong YX, Liang ZJ, et al. Landscape of transcription and expression regulated by DNA methylation related to age of donor and cell passage in adipose-derived mesenchymal stem cells. Aging 2020;12(21):21186–21201; doi: 10.18632/aging.103809.

22. Sandonà M, di Pietro L, Esposito F, et al. Mesenchymal Stromal Cells and Their Secretome: New Therapeutic Perspectives for Skeletal Muscle Regeneration. Front Bioeng Biotechnol 2021;9; doi: 10.3389/fbioe.2021.652970.

23. Coppé JP, Patil CK, Rodier F, et al. Senescence-associated secretory phenotypes reveal cell-nonautonomous functions of oncogenic RAS and the p53 tumor suppressor. PLoS Biol 2008;6(12):2853–68; doi: 10.1371/journal.pbio.0060301.

24. Block TJ, Marinkovic M, Tran ON, et al. Restoring the quantity and quality of elderly human mesenchymal stem cells for autologous cell-based therapies. Stem Cell Res Ther 2017;8(1):239; doi: 10.1186/s13287-017-0688-x.

25. O’Hagan-Wong K, Nadeau S, Carrier-Leclerc A, et al. Increased IL-6 secretion by aged human mesenchymal stromal cells disrupts hematopoietic stem and progenitor cells’ homeostasis. Oncotarget 2016;7(12):13285–96; doi: 10.18632/oncotarget.7690.

26. Gala K, Burdzińska A, Idziak M, et al. Characterization of bone-marrow-derived rat mesenchymal stem cells depending on donor age. Cell Biol Int 2011;35(10):1055–62; doi: 10.1042/cbi20100586.

27. Gnani D, Crippa S, della Volpe L, et al. An early-senescence state in aged mesenchymal stromal cells contributes to hematopoietic stem and progenitor cell clonogenic impairment through the activation of a pro-inflammatory program. Aging Cell 2019;18(3):e12933; doi: 10.1111/acel.12933.

28. Taguchi T, Borjesson DL, Osmond C, et al. Influence of Donor’s Age on Immunomodulatory Properties of Canine Adipose Tissue-Derived Mesenchymal Stem Cells. Stem Cells Dev 2019;28(23):1562–1571; doi: 10.1089/scd.2019.0118.

29. Basisty N, Kale A, Jeon OH, et al. A proteomic atlas of senescence-associated secretomes for aging biomarker development. PLoS Biol 2020;18(1):e3000599; doi: 10.1371/journal.pbio.3000599.

30. Shin S, Lee J, Kwon Y, et al. Comparative proteomic analysis of the mesenchymal stem cells secretome from adipose, bone marrow, placenta and wharton’s jelly. Int J Mol Sci 2021;22(2):845; doi: 10.3390/ijms22020845.

31. Pires AO, Mendes-Pinheiro B, Teixeira FG, et al. Unveiling the Differences of Secretome of Human Bone Marrow Mesenchymal Stem Cells, Adipose Tissue-Derived Stem Cells, and Human Umbilical Cord Perivascular Cells: A Proteomic Analysis. Stem Cells Dev 2016;25(14):1073–83; doi: 10.1089/scd.2016.0048.

32. Wang Z gang, He Z yi, Liang S, et al. Comprehensive proteomic analysis of exosomes derived from human bone marrow, adipose tissue, and umbilical cord mesenchymal stem cells. Stem Cell Res Ther 2020;11(1):511; doi: 10.1186/s13287-020-02032-8.

33. Marote A, Santos D, Mendes-Pinheiro B, et al. Cellular Aging Secretes: a Comparison of Bone-Marrow-Derived and Induced Mesenchymal Stem Cells and Their Secretome Over Long-Term Culture. Stem Cell Rev Rep 2022;19(1):248–263; doi: 10.1007/s12015-022-10453-6.

34. Ratushnyy A, Ezdakova M, Buravkova L. Secretome of senescent adipose-derived mesenchymal stem cells negatively regulates angiogenesis. Int J Mol Sci 2020;21(5):1802; doi: 10.3390/ijms21051802.

35. Chou LY, Ho C te, Hung SC. Paracrine Senescence of Mesenchymal Stromal Cells Involves Inflammatory Cytokines and the NF-κB Pathway. Cells 2022;11(20):3324; doi: 10.3390/cells11203324.

36. Severino V, Alessio N, Farina A, et al. Insulin-like growth factor binding proteins 4 and 7 released by senescent cells promote premature senescence in mesenchymal stem cells. Cell Death Dis 2013;4(11):e911; doi: 10.1038/cddis.2013.445.

37. Salerno A, Brady K, Rikkers M, et al. MMP13 and TIMP1 are functional markers for two different potential modes of action by mesenchymal stem/stromal cells when treating osteoarthritis. Stem Cells 2020;38(11):1438–1453; doi: 10.1002/stem.3255.

38. Siennicka K, Zołocińska A, Dȩbski T, et al. Comparison of the Donor Age-Dependent and in Vitro Culture-Dependent Mesenchymal Stem Cell Aging in Rat Model. Stem Cells Int 2021;2021:6665358; doi: 10.1155/2021/6665358.

39. Taylor SE, Clegg PD. Collection and Propagation Methods for Mesenchymal Stromal Cells. Veterinary Clinics of North America - Equine Practice 2011;27(2):263–274; doi: 10.1016/j.cveq.2011.05.003.

40. Caffi V, Espinosa G, Gajardo G, et al. Pre-conditioning of Equine Bone Marrow-Derived Mesenchymal Stromal Cells Increases Their Immunomodulatory Capacity. Front Vet Sci 2020;7; doi: 10.3389/fvets.2020.00318.

41. Gale AL, Linardi RL, McClung G, et al. Comparison of the chondrogenic differentiation potential of equine synovial membrane-derived and bone marrow-derived mesenchymal stem cells. Front Vet Sci 2019;6:178; doi: 10.3389/fvets.2019.00178.

42. Ca S, Ws R, Kw E. NIH Image to ImageJ: 25 years of image analysis. Nat Methods 2012;9(7):671–675.

43. Bundgaard L, Stensballe A, Elbæk KJ, et al. Mapping of equine mesenchymal stromal cell surface proteomes for identification of specific markers using proteomics and gene expression analysis: An in vitro cross-sectional study. Stem Cell Res Ther 2018;9(1):288; doi: 10.1186/s13287-018-1041-8.

44. Schmittgen TD, Livak KJ. Analyzing real-time PCR data by the comparative CT method. Nat Protoc 2008;3(6):1101–8; doi: 10.1038/nprot.2008.73.

45. Hammond DE, Kumar JD, Raymond L, et al. Stable isotope dynamic labeling of secretomes (SIDLs) identifies authentic secretory proteins released by cancer and stromal cells. Molecular and Cellular Proteomics 2018;17(9):1837–1849; doi: 10.1074/mcp.TIR117.000516.

46. Cox J, Hein MY, Luber CA, et al. Accurate proteome-wide label-free quantification by delayed normalization and maximal peptide ratio extraction, termed MaxLFQ. Molecular and Cellular Proteomics 2014;13(9):2513–2526; doi: 10.1074/mcp.M113.031591.

47. Tyanova S, Temu T, Cox J. The MaxQuant computational platform for mass spectrometry-based shotgun proteomics. Nat Protoc 2016;11(12):2301–2319; doi: 10.1038/nprot.2016.136.

48. Cox J, Neuhauser N, Michalski A, et al. Andromeda: A peptide search engine integrated into the MaxQuant environment. J Proteome Res 2011;10(4):1794–1805; doi: 10.1021/pr101065j.

49. Yu G, Wang LG, Han Y, et al. ClusterProfiler: An R package for comparing biological themes among gene clusters. OMICS 2012;16(5):284–7; doi: 10.1089/omi.2011.0118.

50. Wu T, Hu E, Xu S, et al. clusterProfiler 4.0: A universal enrichment tool for interpreting omics data. The Innovation 2021;2(3):100141; doi: 10.1016/j.xinn.2021.100141.

51. Sardá-Espinosa A. Time-series clustering in R Using the dtwclust package. R Journal 2019;11(1); doi: 10.32614/rj-2019-023.

52. Kumar L, Futschik ME. Mfuzz: A software package for soft clustering of microarray data. Bioinformation 2007;2(1):5–7; doi: 10.6026/97320630002005.

53. Gaujoux R, Seoighe C. A flexible R package for nonnegative matrix factorization. BMC Bioinformatics 2010;11:367; doi: 10.1186/1471-2105-11-367.

54. Ziegler J, Vogt T, Miersch O, et al. Concentration of dilute protein solutions prior to sodium dodecyl sulfate polyacrylamide gel electrophoresis. Anal Biochem 1997;250(2):257–60; doi: 10.1006/abio.1997.2248.

55. Marx C, Gardner S, Harman RM, et al. The mesenchymal stromal cell secretome impairs methicillin-resistant Staphylococcus aureus biofilms via cysteine protease activity in the equine model. Stem Cells Transl Med 2020;9(7):746–757; doi: 10.1002/sctm.19-0333.

56. Marinkovic M, Dai Q, Gonzalez AO, et al. Matrix-bound Cyr61/CCN1 is required to retain the properties of the bone marrow mesenchymal stem cell niche but is depleted with aging. Matrix Biology 2022;111:108–132; doi: 10.1016/j.matbio.2022.06.004.

57. LeClair RJ, Durmus T, Wang Q, et al. Cthrc1 is a novel inhibitor of transforming growth factor-β signaling and neointimal lesion formation. Circ Res 2007;100(6):826–33; doi: 10.1161/01.RES.0000260806.99307.72.

58. Duan X, Yuan X, Yao B, et al. The Role of CTHRC1 in Promotion of Cutaneous Wound Healing. Signal Transduct Target Ther 2022;7(1); doi: 10.1038/s41392-022-01008-9.

59. Ruiz-Villalba A, Romero JP, Hernández SC, et al. Single-Cell RNA Sequencing Analysis Reveals a Crucial Role for CTHRC1 (Collagen Triple Helix Repeat Containing 1) Cardiac Fibroblasts After Myocardial Infarction. Circulation 2020;142(19); doi: 10.1161/CIRCULATIONAHA.119.044557.

60. Kagan HM, Li W. Lysyl oxidase: Properties, specificity, and biological roles inside and outside of the cell. J Cell Biochem 2003;88(4):660–72; doi: 10.1002/jcb.10413.

61. Rozanova S, Barkovits K, Nikolov M, et al. Quantitative Mass Spectrometry-Based Proteomics: An Overview. In: Methods in Molecular Biology 2021; pp. 86–116; doi: 10.1007/978-1-0716-1024-4_8.

62. Arnhold S, Elashry MI, Klymiuk MC, et al. Investigation of stemness and multipotency of equine adipose-derived mesenchymal stem cells (ASCs) from different fat sources in comparison with lipoma. Stem Cell Res Ther 2019;10(1):309; doi: 10.1186/s13287-019-1429-0.

63. Song EK, Jeon J, Jang DG, et al. ITGBL1 modulates integrin activity to promote cartilage formation and protect against arthritis. Sci Transl Med 2018;10(462):eaam7486; doi: 10.1126/scitranslmed.aam7486.

64. Braga CL, da Silva LR, Santos RT, et al. Proteomics profile of mesenchymal stromal cells and extracellular vesicles in normoxic and hypoxic conditions. Cytotherapy 2022; doi: 10.1016/j.jcyt.2022.08.009.

65. Li XQ, Zhang R, Lu H, et al. Extracellular Vesicle-Packaged CDH11 and ITGA5 Induce the Premetastatic Niche for Bone Colonization of Breast Cancer Cells. Cancer Res 2022;82(8):1560–1574; doi: 10.1158/0008-5472.CAN-21-1331.

66. Mathieu M, Névo N, Jouve M, et al. Specificities of exosome versus small ectosome secretion revealed by live intracellular tracking of CD63 and CD9. Nat Commun 2021;12(1); doi: 10.1038/s41467-021-24384-2.

67. Won E, Morodomi Y, Kanaji S, et al. Extracellular tyrosyl-tRNA synthetase cleaved by plasma proteinases and stored in platelet α-granules: Potential role in monocyte activation. Res Pract Thromb Haemost 2020;4(7); doi: 10.1002/rth2.12429.

