## Supplementary File 1 for "Mesenchymal Stromal Cell secretome is affected by tissue source, donor age and sex"

### Isolation of adipose-derived mesenchymal stromal cells (ASCs)

Adipose tissue collected during abdominal surgery was minced with surgical blade, washed with PBS and digested in equal volume of PBS with 1% bovine serum albumin an 0.1% collagenase type I, by placing on orbital shaker for 50min at 37◦C. Following digestion, solution was filtered through 70μm cell strainer and centrifuged at 260*g* for 5min. Resulting cell pellet was washed in PBS at 300*g* for 5min and seeded in T75 flask, where they were grown until 70% confluence, passaged and cryopreserved.

### Tri-lineage differentiation assay

Cells were seeded in 12-well plates in duplicates for osteogenic (5,000 cells/$cm^2$) and adipogenic (10,000 cells/$cm^2$) differentiation assays. After 48h, differentiation was induced in one of the replicates with StemPro Osteogenesis and Adipogenesis differentiation kits respectively, following the manufacturer's instructions. The other replicate was maintained in the expansion medium and served as negative control. For chondrogenic differentiation, cells were cultivated in duplicates pellet culture[1]. Briefly, 0,5 x $10^6$ cells were suspended in conical microtube and centrifuged at 400g for 10min. After 48h, chondrogenesis was induced in one replicate with chondrogenic media (StemPro Chondrogenesis Differentiation Kit) while the other pellet was maintained in the expansion medium. Following 14 days of culture for osteogenic and adipogenic assays and 21 days for chondrogenic assay, cells were washed in PBS and fixed in 10% neutral buffered formalin prior to staining.

Pellet cultures were transferred to a 96-well plate and microscopic image taken before paraffin embedding and sectioning. Two percent Alizarin red solution (Sigma-Aldrich) was used to identify calcium deposits in osteogenic culture, 0,5% solution of Oil Red O in isopropanol (Sigma-Aldrich) was used to identify lipid droplets in adipogenic culture and 1% Alcian Blue solution (Sigma-Aldrich) was used to detect glycosaminoglycans in cell pellet sections in chondrogenic assay.

### Protein digestion for LC-MS/MS

Protein-bound StrataClean resin was re-suspended in 160 µl of 25 mM ammonium bicarbonate (ambic, Fluka Chemicals, UK). To enhance protein solubilisation, 1% (w/v) RapiGest (Waters, UK) in 25 mM ambic was added and sample heated at 80 °C for 10 min. Proteins were reduced with dithiothreitol (3 mM final, Sigma-Aldrich) for 10 min at 60 °C, followed by alkylation for 30 min in the dark using iodoacetamide (9 mM final, Sigma-Aldrich). For the first protease digestion, trypsin/LysC (MS grade trypsin/LysC Mix, Promega) was added and samples incubated at 37 °C for 16 h on a rotary mixer, followed by second digestion at 37 °C for 2 h. Trypsin/Lys-C were used at an enzyme to substrate ratio of 1:50 (w/w). Peptide digests were acidified with trifluoroacetic acid (TFA, Sigma-Aldrich) at a final concentration of 0.5% (v/v) and incubated at 37 °C for 45 min. Samples were centrifuged at 16,000 x g for 15 min, supernatants separated from the StrataClean resin and centrifuged again for 10 min to remove any residual insoluble material.

### LC-MS/MS spectral acquisition

The sample was loaded onto the trapping column (Thermo Scientific, PepMap100, C18, 300 µm x 5 mm), using partial loop injection, for seven minutes at a flow rate of 12 µL/min with 0.1% (v/v) FA. The sample was resolved on the analytical column (Easy-Spray C18 75 µm x 500 mm 2 µm column) using a gradient of 96.2% A (0.1% formic acid) 3.8% B (79.95% acetonitrile, 19.95% water, 0.1% formic acid) to 50% A 50% B over 90 minutes at a flow rate of 0.3 nL/min (2-hour gradient), interspersed with

30min blanks between the samples. The data-dependent program used for data acquisition consisted of a 60,000-resolution full-scan MS scan in the orbitrap (AGC set to 3e6 ions with a maximum fill time of 100ms). The 16 most abundant peaks per full scan were selected for HCD MS/MS (30,000 resolution, AGC set to 1e5 ions with a maximum fill time of 45 ms) with an ion selection window of 2 m/z and normalised collision energy of 30 %. Ion selection excluded singularly charged ions and ions with equal to or a greater than +6 charge state. To avoid repeated selection of peptides for fragmentation the program used a 60-second dynamic exclusion window. All samples were analysed in random order.

**Peptide identification and quantification**

Protein identification with MaxQuant software was based on minimum one peptide, the minimum peptide length of seven amino acids, trypsin/P as proteolytic enzyme, single missed cleavage, 'match between runs' function enabled. Mass deviations were set to 20 ppm on the precursor ion and 0.5 Da on the fragment ions. Cysteine carbamidomethylation was set as a fixed modification. Methionine oxidation and acetylation at the protein N-terminus was allowed as a variable modification.

**Imputation methods**

The presence of missing values in proteomics data, particularly those seen missing not at random, can lead to a skewed distribution, impacting on downstream analysis results. The use of imputation methods is commonly used to improve proteome coverage and therefore improve statistical power to the analysis[2].

We compared the effects of handling the missing values using three different methods; i) using no imputation, ii) using sample-based imputation methods and iii) using protein based imputation methods. For the no imputation method, the data was filtered for 30% "missingness" and the remaining missing values were carried forward as "N/A" missing values. For the sample-based imputation methods, we looked at the proportion of missing values per sample. Samples with more than 10% of proteins missing were considered to be missing not at random. Samples with less than 10% of proteins showing missing values were assumed to be missing at random. For the protein-based imputation, samples were grouped by condition, ie. tissue type, and proteins with more than 30% missing values within a condition were assumed to be linked to sample condition and therefore missing not at random. All other missing values were assumed missing at random. For both the sample and protein-based imputation methods, missing not at random values were assigned values of zero. Missing at random values were imputed using the knn (nearest neighbour) method using the DEP Bioconductor R package[3,4].

In order to determine the effect of missing values on our results, we compared the effects of using these imputation methods within our analysis. It was found that the different methods of handling the missing values within our data resulted in similar clustering. Comparing the PCA analysis of each method (Supplementary Figure 1), little difference can be seen between the imputation methods. Similarly, the handling of missing values showed very little impact on the time course clustering methods (data not shown). The imputation methods therefore weren't shown to add strength or statistical power to our analysis. As a result, it was decided to report the results of the analysis using the non-imputed values, i.e. true values, and these results were used for the entire analysis.

**Lysyl oxidase activity assay**

Lysyl oxidase activity was detected as fluorescence at Ex/Em = 544/590 nm after 10, 20 and 30 minutes of sample incubation with proprietary substrate using FluorSTAR Optima microplate reader (BMG Labtech). Fluorescence intensity was expressed relative to total protein concentration of the sample.

To test hypothesis that activity of lysyl oxidase (LOX) in MSC CM decreases with donor age, we analysed change in fluorescence over time in samples incubated with proprietary substrate, leading to release of hydrogen peroxide detected in horseradish peroxidase-coupled reaction (Lysyl Oxidase Activity Assay Kit, Abcam). Each sample was analysed in duplicate. Results recorded as relative fluorescence units (RFU) were blank corrected and divided by total protein concentration of the sample. Normalised fluorescence values at 10, 20 and 30 minutes of CM incubation were used to fit generalised linear mixed-effect model with Gamma distribution using glmer() function in the package lme4 (v.1.1.31) in R. Age and time variables were scaled and centred by subtracting the mean and dividing by standard deviation of each variable prior to fitting the model. Age, time and their interaction were included as fixed effects and donor as random effect. The effect of predictors was considered significant when t-statistics correspond to $p<0.05$.

**Western Blot analysis of collagen triple helix repeat-containing protein 1 (CTHRC1)**

All reagents were sourced from ThermoFisher Scientific unless stated otherwise. Samples were concentrated using StrataClean resin as described previously[54], using 10µl of StrataClean and the CM volume adequate to obtain 20µg resin-bound protein in each sample. Samples were denatured and reduced through heating for 10 min at 70°C with NuPAGE Sample Reducing Agent and Novex Tris-Glycine SDS Sample Buffer. StrataClean resin was separated through centrifugation at 295g for 1 min and supernatant applied directly to SDS polyacrylamide gel electrophoresis (NuPage 4-12% Bis-Tris Mini Gel). Protein standard (1:1 mix of Novex Sharp Pre-Stained and MagicMark™ XP standards) was loaded along the samples for molecular weight estimation. Protein was transferred to nitrocellulose membrane using XCell system for 60 min at 30V and membrane blocked for 30 min with StartingBlock (TBS). After blocking, membrane was stained with rabbit anti-human CTHRC1 antibody (Abcam, ab85739) diluted 1:500 in StartingBlock. Primary antibody incubation was undertaken on a rotating platform at 4°C overnight. Membranes were washed three times for 5 min in tris-buffered saline (TBS) and incubated with goat anti-Rabbit fluorescently labelled secondary antibody (WesternDot 625) diluted 1:500 in StartingBlock for 1 hour at room temperature. Membranes were washed three times for 5 min in TBS and imaged with UV transilluminator (XXXXXX) and UviPro software (XXXX). Fibronectin detected with anti-human antibody (Sigma-Aldrich, F3648) diluted 1:1000 was used as a loading control[55].

Relative quantification of CTHRC1 abundance was performed using ImageJ. Mean grey value was measured for each CTHRC1 and FBN band, and their backgrounds, using region of interest (ROI) standardised for each protein. Pixel density measurements have been inverted and background value was subtracted from each band value. Relative density of CTHRC1 was then calculated as CTHRC1/FBN density ratio.

**ELISA analysis of monocyte chemoattractant protein 1 (MCP-1)**

MCP-1 concentration was measured in 24 samples of MSC CM (13 ASC and 11 BMSC) in duplicate based on the absorbance measured at 450nm using Multiskan FC microplate reader (Thermo Scientific). MCP-1 concentration was calculated based on the blank corrected absorbance read using the standard curve plotted on the log-log axes. Concentration value in one sample has been considered an outlier based on the threshold of three standard deviations of the mean and removed from the dataset. Samples with MCP-1 concentrations below the assay detection threshold (8.19

pg/ml) were imputed to arbitrary value below the limit of detection (8.1 pg/ml). Next, we used GLM with Gamma distribution and log link to test association between MCP-1 concentration and MSC donor age, as described for mass spectrometry data. The results were robust to other tested assumptions (removal of samples below detection threshold from the dataset, imputation at 4 pg/ml) and therefore it was decided to report results with the values imputed at the lower limit of detection of the assay.

**Stable isotope labelling identified secreted proteins in MSC CM based on their kinetic behaviour**

In this dataset, 163 proteins met the criteria required for RIA trajectory analysis over time (peptides associated with the protein quantified in at least three time points) and 108 proteins were characterised with k value. The remaining 55 proteins were unfit for analysis due to RIA assigned to their peptides assuming only values of 0 or 1, which was related with detection of only 'light' or 'heavy' form of the peptide. Out of 108 proteins, 106 were characterised with flux and 2 remaining proteins were excluded due to negative P value (mean total abundance higher at 6 h than 24 h time point).
